# A possible role for epigenetic feedback regulation in the dynamics of the Epithelial-Mesenchymal Transition (EMT)

**DOI:** 10.1101/651620

**Authors:** Wen Jia, Abhijeet Deshmukh, Sendurai A. Mani, Mohit Kumar Jolly, Herbert Levine

**Affiliations:** Center for Theoretical Biological Physics, Rice University, Houston, TX 77005, USA; Department of Physics and Astronomy, Rice University, Houston, TX 77005, USA; Department of Translational Molecular Pathology, University of Texas MD Anderson Cancer Center, Houston, TX 77005, USA; Centre for BioSystems Science and Engineering, Indian Institute of Science, Bangalore 560012, India; Department of Bioengineering, Rice University, Houston, TX 77005, USA; Department of Bioengineering, Northeastern University, Boston, MA 02115, US; Department of Physics, Northeastern University, Boston, MA 02115, USA

**Keywords:** Epithelial-Mesenchymal Transition, Mathematical modeling, Epigenetic regulation

## Abstract

The epithelial-mesenchymal transition (EMT) often plays a critical role in cancer metastasis and chemoresistance, and decoding its dynamics is crucial to design effective therapeutics. EMT is regulated at multiple levels – transcriptional, translational, protein stability, and epigenetics; the mechanisms by which epigenetic regulation can alter the dynamics of EMT remain elusive. Here, to identify the possible effects of epigenetic regulation in EMT, we incorporate a feedback term in our previously proposed model of EMT regulation of the miR-200/ZEB/miR-34/SNAIL circuit. This epigenetic feedback that stabilizes long-term transcriptional activity can alter the relative stability and distribution of states in a given cell population, particularly when incorporated in the inhibitory effect on miR-200 from ZEB. This feedback can stabilize the mesenchymal state, thus making transitions out of that state difficult. Conversely, epigenetic regulation of the self-activation of ZEB has only minor effects. Our model predicts that this effect could be seen in experiments, when epithelial cells are treated with an external EMT-inducing signal for a sufficiently long period of time and then allowed to recover. Our preliminary experimental data indeed shows that a prolonged TGF-β exposure gives rise to increasing difficult reversion back to the epithelial state. Thus, this integrated theoretical-experimental approach yields insights into how an epigenetic feedback may alter the dynamics of EMT.

## Introduction

Cancer metastasis refers to the spreading of cancer cells from primary tumor to different parts of the body and the forming of secondary tumors. It leads to more than 90% of cancer-related deaths (1). Extensive research has identified the role of partial/complete EMT (Epithelial-Mesenchymal Transition) and its reverse MET (Mesenchymal-Epithelial Transition) in forming metastases (2,3). Thus, understanding the detailed dynamics of EMT and MET is crucial for decoding and halting metastases.

EMT refers to a trans-differentiation process whereby cells change their phenotype from being primarily epithelial (tight cell-cell adhesion, apico-basal polarity, limited invasion and migration) to having traits typical of mesenchymal lineage(little/no cell-cell adhesion, front-back polarity, having increased invasion and migration capabilities). Epithelial cells express high level of E-cadherin (encoded by the CDH1 gene), a cell-cell adhesion molecule that often can be used as an epithelial marker. On the contrary, the intermediate filament Vimentin is a typical mesenchymal marker (4). Cancer cells undergoing EMT have increased chance of leaving the primary tumor and transiting through the circulation. It is then hypothesized that these cells undergo the reverse process (MET) to regain their cell-cell adhesion and form a new tumor, once they reach a distant organ (4). Recent studies have indicated that cells can exist in intermediate states that exhibit both epithelial and mesenchymal markers and traits – hybrid E/M phenotypes (5,6,7,8) – which may enhance collective cell migration, leading to the formation of clusters of circulating tumor cells (CTCs) that can be highly tumorigenic (9).

A previously proposed and validated dynamical model of the core EMT regulatory circuit is based on two mutually inhibiting microRNA-TF loops: miR-200/ZEB and miR-34/SNAIL (10). ZEB and SNAIL are two transcription factors (TFs) that repress the transcription of miR-34 and miR-200, while miR-34 and miR-200 can downregulate the protein production of ZEB and SNAIl through binding with the corresponding mRNA and inhibiting its translation and/or increasing its degradation. High SNAIL and ZEB levels correspond to a mesenchymal phenotype, and high miR-200/miR-34 levels correspond to an epithelial phenotype (11,12,13,14). This model predicted the existence of a stable hybrid epithelial/ mesenchymal (E/M) phenotype that has subsequently been observed *in vitro* (15) and *in vivo* (16,17). Further, this model predicted hysteresis or ‘cellular memory’ in EMT/MET dynamics that has been observed for multiple cell lines undergoing EMT in response to TGFβ (transforming growth factor β) (18,19). However, the full set of possible mechanisms underlying hysteresis in EMT remains elusive.

Epigenetic mechanisms have been proposed to give rise to cellular memory in multiple contexts (20, 21, 22, 23); however, the abovementioned as well as other models of EMT regulatory networks do not consider the crucial role of epigenetics (24). Epigenetics refers to mechanisms that can hertitably alter cellular phenotypes without changing the genome or nucleotide sequence (25). Epigenetic regulation can be achieved in (at least) two different ways - DNA methylation or modification of histone proteins. DNA wraps around histone proteins forming nucleosomes and epigenetic modification of the histone protein “tails” can affect the extent of which nucleosomes are compacted. These factors can, in turn, affect the gene expression by altering the chromatin structure to create transcriptionally more active or more inactive regions (26, 27). Some studies have reported how epigenetic silencing of E-cadherin can emerge after an EMT process (25). Also, specific histone acetylation or demethylation can suppress the induction as well as the maintenance of SNAIL-1-mediated EMT (28, 29). However, whether epigenetics can give rise to ‘cellular memory’ in the context of EMT has not been well-studied.

Epigenetic modifications are known to regulate cell differentiation during development, where specific chromatin-based mechanisms have been proposed to maintain pluripotency (30). The exact molecular mechanisms by which this epigenetic regulation occurs are complex and multi-faceted; hence there is no agreed-upon methodology for simulating the role of this epigenetic feedback. A phenomenological model to investigate the epigenetic process during cell differentiation introduced variables that can modulate the thresholds for the influences of transcription factors on gene expression of their downstream targets (31). The underlying idea in this framework is that a given gene expression state can be stabilized by epigenetic feedback regulation, i.e. the longer a gene is turned on, the easier it becomes for it to stay transcriptionally active, thus giving rise to extremely hysteretic dynamics.

Here, we use this conceptual framework to study the possible role of epigenetic feedback in EMT. Specifically, we incorporate similar phenomenological terms into our previously developed dynamical model of EMT and analyze the effects of expression-dependent modulation of several threshold parameters in the model, and the consequences for phenotypic switching dynamics. Our results demonstrate that epigenetic feedback acting on the inhibition on miR-200 by ZEB would significantly stabilize the mesenchymal state, thus making EMT irreversible in some cells, while a similar feedback acting on the self-activation of ZEB barely has any major consequence on EMT dynamics. The model allows us to simulate a specific experimental scenario involving prolonged TGF-β exposure to epithelial cells followed by attempted recovery; as expected, a prolonged exposure enables epigenetic feedback to dramatically change transition rates, leading to effective irreversibility of EMT. This prediction is confirmed by preliminary experimental data in MCF10A cells with live-cell dual imaging reporters that can measure epithelial and mesenchymal markers simultaneously. Thus, our integrated computational-experimental approach highlights the potential importance of epigenetic feedback in governing EMT dynamics.

## Results

### Epigenetic feedback on self-activation of ZEB does not largely affect the EMT dynamics

In this study, we incorporate epigenetic feedback into our previously developed model of a core EMT/MET regulatory network---the miR-200/ZEB and the miR-34/SNAIL mutually inhibiting loops (Fig. 1A; SI sections 1 and 2). Various environmental signals can activate SNAIL that can then drive the miR-200/ZEB switch towards the mesenchymal state. The miR-200/ZEB loop can behave as a tristable circuit, enabling three states – epithelial (high miR-200, low ZEB), mesenchymal (low miR-200, high ZEB) and hybrid epithelial/mesenchymal (medium miR-200, medium ZEB) (Fig. 1B) (10). Thus, the two main determinants of the state of this switch, in terms of threshold levels, are the thresholds for transcriptional regulation by ZEB, either for the inhibition on miR-200 or its self-activation (possibly occurring indirectly through other factors such as ESRP1) (32).

**Figure 1.**
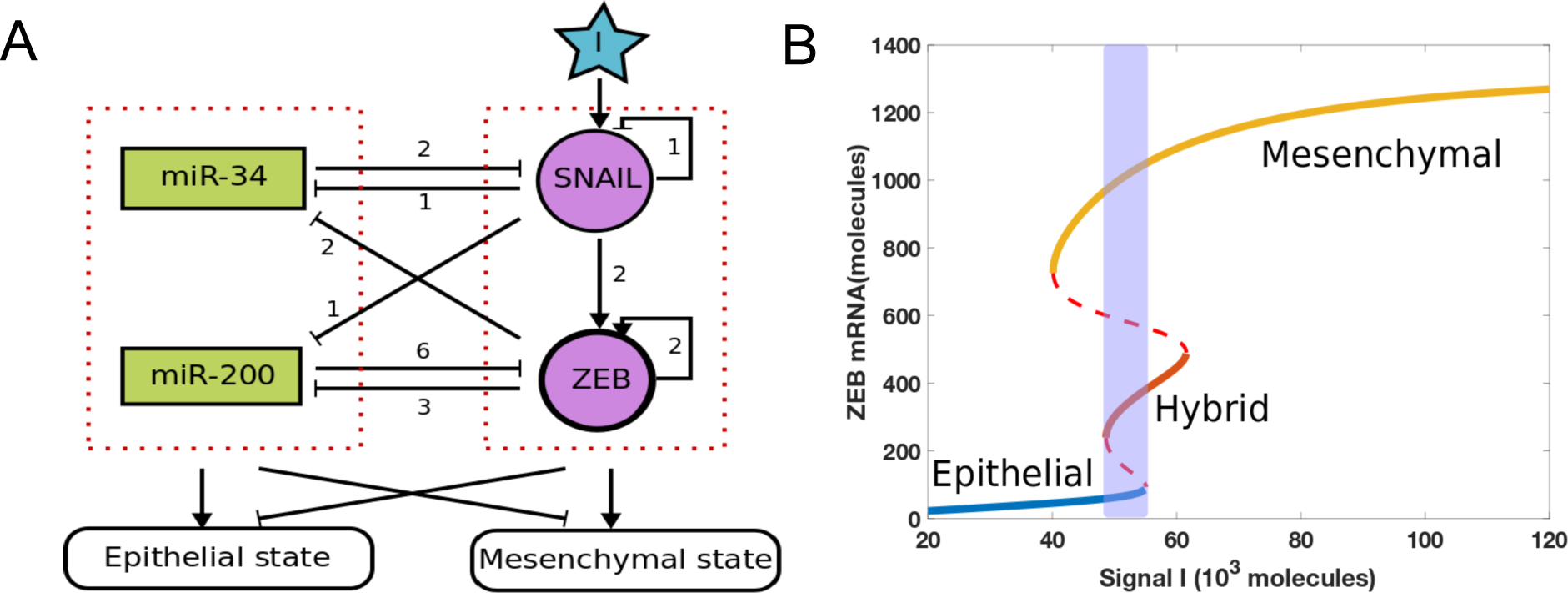
(A) A core regulatory network for EMT that consists of two highly linked microRNA-TF mutually inhibitory circuits : miR-34/SNAIL and miR-200/ZEB. Signal I represents external signals such as HGF, NF-*κ*B, Wnt, TGF-*β* and/or HIF1*α*. The numbers represent the number of binding sites obtained from experiments for these different biological processes (10). (B) Bifurcation diagram of ZEB mRNA levels for the network shown in Fig 1A, with I as the bifurcation parameter. Solid lines represent stable states, dashed lines represent unstable states. The shaded region shows a tristable parameter range.

The bifurcation diagram for this circuit shows monostable and multistable phases, including a tristable region indicating the coexistence of epithelial (E; ZEB mRNA <100 molecules), hybrid E/M (E/M; 240 <ZEB mRNA< 490 molecules) and mesenchymal (M; ZEB mRNA >730 molecules) (Fig. 1B). In the presence of stochastic effects due to transcriptional noise, individual cells can transition between the various states present at a fixed value of the external signal, leading eventually to a mixed population with different percentages of the three phenotypes. To illustrate this, we incorporated noise (exponentially correlated in time) to an external signal I driving the governing equations for this circuit and simulated the resultant stochastic dynamics of a population of 1000 cells (SI section 3). The (mean) value of the signal I is fixed at the midpoint of the tristable region, and all modeled cells are in a stable E state initially. We observed a stable distribution at around time 20 *ζ* with 75% E, 15% E/M and 10% M (dashed lines in Fig 2B). This distribution serves as a baseline for considering the effects of adding an epigenetic feedback. The time unit *ζ* = 100 hours here is the time factor used in the epigenetic feedback term (see SI section 4).

**Figure 2.**
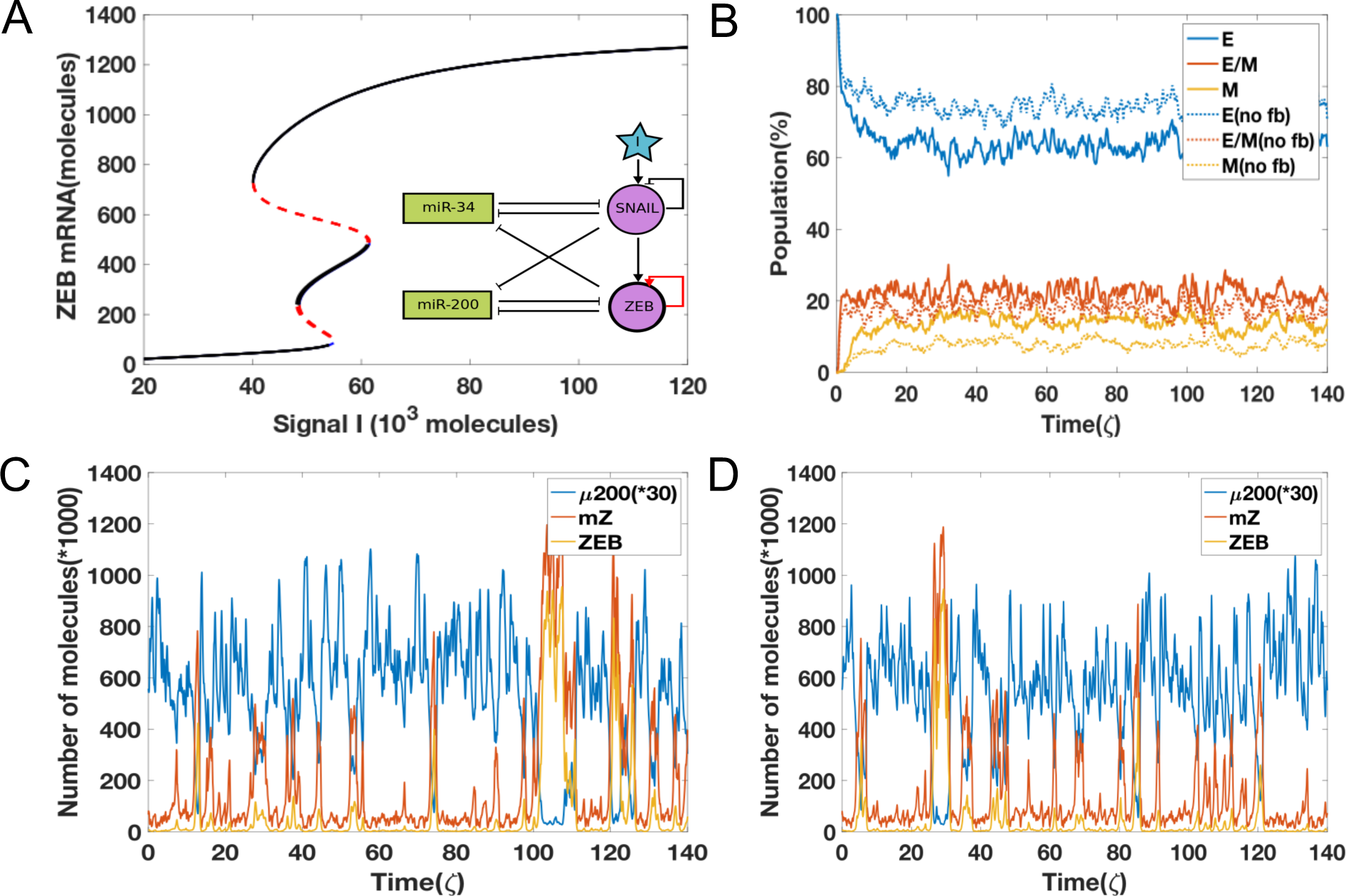
(A) The bifurcation diagrams for the EMT circuit shown in Fig 1, for varying values of *α* when epigenetic feedback acts through ZEB self-activation. The black curves representing stable states at finite feedback almost overlap with the blue ones for the no feedback case; red lines denote unstable states. (B) Simulations showing the population change as a function of time. The percentage is calculated based on 1000 independent simulations. Dashed lines represent the no epigenetic feedback case (*α* = 0), and solid lines are with feedback (*α* = 0.017). (C) A sample dynamic plot for the no feedback case. (D) A sample dynamic plot with the inclusion of feedback.

Next, we added strong epigenetic feedback governing the threshold which controls the self-activation of ZEB. Specifically, the threshold now obeys a dynamical equation that lowers its steady-state value by an amount proportional to the modulation factor α in a state with high ZEB (SI section 4). Adding this feedback barely changes the bifurcation result (Fig. 2A); this is true for all reasonable values of α (SI section 4). Furthermore, this feedback also barely changes the population structure (solid lines in Fig. 2B) – the stable distribution now is 65% E, 20% E/M, and 15% M. We also see minimal differences in the dynamics of individual cells in the case of no feedback (Fig. 2C) vs. that with epigenetic feedback (Fig. 2D); in both cases, we see both partial EMT/MET as well as complete EMT/MET. Together, these results convey that the epigenetic feedback acting on self-activation of ZEB can only slightly change the dynamic properties of EMT. Essentially, a state that has high ZEB (namely M) is not sensitive to the fact that the threshold has been lowered and transitions out of this state will mostly involve fluctuations that increase miR-200 and its strong inhibition, and will again be relatively insensitive to the ZEB self-activation. Similar observations are seen in a simpler case with similar network topology – a SATS (self-activated toggle switch) that does not incorporate microRNA-medidated dynamics (Fig. S1, S2) (33).

### Epigenetic feedback on the inhibition of miR-200 by ZEB can stabilize a mesenchymal state

Based on results from the SATS model (SI section 5, 6, 7), we expect that modifying the transcriptional inhibition of miR-200 by ZEB would have a more pronounced effect in terms of population distribution. Indeed, unlike the previous results, increasing the epigenetic feedback on the inhibition of miR-200 by ZEB significantly alters the bifurcation diagram; now, achieving a mesenchymal state becomes possible at lower values of I (Fig. 3A). Thus, this epigenetic feedback can not only induce EMT, but also potentially increase the stability of the mesenchymal state as hysteresis becomes more prominent. The equilibrium population distribution is also drastically affected; the larger the modulation parameter *α*, the higher the change in the frequencies of different subpopulations in the equilibrium distribution (Fig. 3B). Consistent with the bifurcation diagram, including the epigenetic feedback increases the mesenchymal population at the expense of the epithelial population. Thus, once the epigenetic feedback is strong enough, the change in population distribution can be dramatic. Moreover, it takes much longer for the system to reach a steady state in terms of population distribution, in the presence of a strong epigenetic feedback – without feedback, the system reaches steady state in around time 20 *ζ* (Fig. 3C), but with feedback, this timescale is extended to 50 *ζ* (Fig 3D).

**Figure 3.**
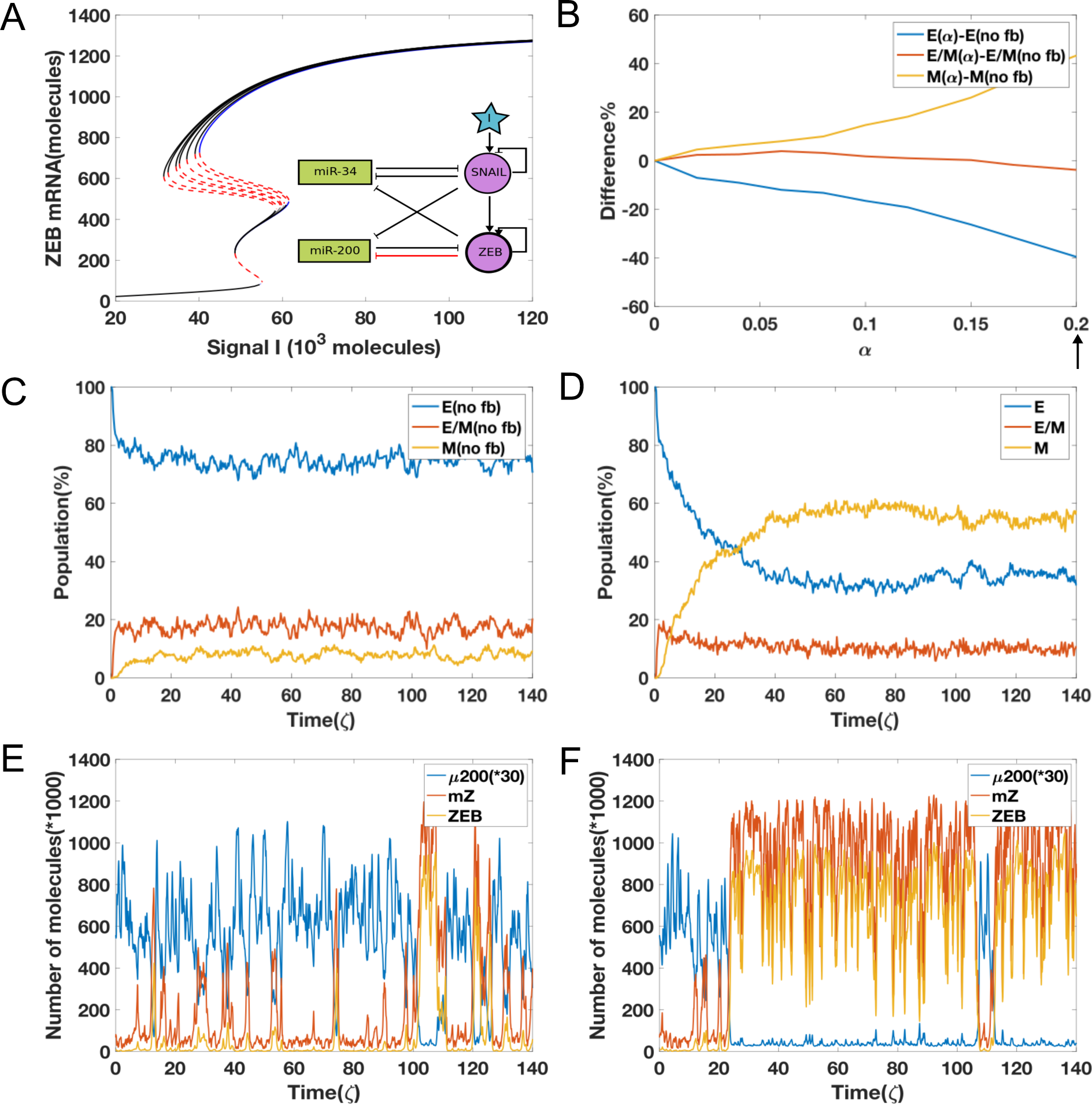
(A) The bifurcation diagrams for the EMT circuit shown in Fig 1A, for varying values of *α* when epigenetic feedback acts through the inhibition on miR-200 by ZEB. With increasing feedback, the bifurcation moves toward the left side in the mesenchymal region. The blue curve represents stable states in no feedback case and the black lines denote stable states with feedback; red lines denote unstable states. (B) The difference between equilibrium distribution population as a function of *α*. (C) A sample plot for the evolution of equilibrium population distribution for 1000 cells, as a function of time without any feedback (*α* = 0). (D) Same as Fig 3C but with strong feedback (*α* value is marked by arrow in 3(B), *α* = 0.2). (E) A sample dynamics plot for without feedback case (same as Fig 2C). (F) A sample dynamic plot for the case with feedback (*α* = 0.2).

The reason for this delayed behavior can be seen by comparing individual simulation runs for the two cases. For the case without any epigenetic feedback, the cell transitions quickly from all of three different states to one another (Fig. 3E). Thus, it is possible for cells that have undergone EMT to revert. In contrast, for the case with a strong epigenetic feedback, cells not only can go through a complete EMT process but also they can become trapped in the mesenchymal state for a much longer period (Fig. 3F). This effect is qualitatively seen for varying strengths of external noise, although the precise time scales seen in our simulation also strongly depends on the strength of the external noise (SI section 3, 9). These results indicate that unlike what we saw previously by modulating the self-activation of ZEB, incorporating an epigenetic feedback into the dynamics of inhibition on miR-200 by ZEB significantly changes the population distribution and stabilizes a mesenchymal phenotype.

Next, we investigated whether a more stable mesenchymal state can render some cells ‘locked’ in that state, i.e. make the transition effectively irreversible. To quantify the EMT irreversibility as caused by this epigenetic regulation, cells are exposed to an external EMT-inducing signal such as TGF-β for varying durations of treatment, and then the signal is withdrawn. If TGF-β is provided continuously (i.e. no withdrawal), the entire population turns mesenchymal (34). But if TGF-β is removed after some time, mesenchymal cells can undergo MET and go back to the previous epithelial or a hybrid E/M phenotype (34). The kinetics of this reversal can be dramatically altered by the epigenetic feedback.

Starting with all cells in an epithelial state, when the external signal I corresponds to the bistable {E, M} part of the bifurcation diagram, we increased the value I (to mimic the treatment with TGF-β) to that in a bistable region {E/M, M} and then decreased it back to its original value (to mimic the action of removing TGF-β) which corresponds to the bistable region {E, M}. Depending on the time for which cells were treated with TGF-β, the population response can vary. If the level of I is reduced very early, most cells immediately revert to being epithelial (Fig. 4A). When the duration of treatment of high I levels increases, some cells maintain a mesenchymal state even after reducing the levels of I (Fig. 4B, 4C). Also, the time for reversal to the initial condition increases as the time duration of treatment with high I becomes longer. If the duration of high I treatment increases to around time 9.6 *ζ*, all cells are able to stay in mesenchymal state (Fig. 4D). This effect can thus alter the population distribution of cells exposed to signal I for varying duration; most cells tend to acquire an irreversible transition around time 5-7 *ζ*. After 9 *ζ* of exposure to high I, all cells can be ‘locked’ in a mesenchymal state (Fig. 5).

**Figure 4.**
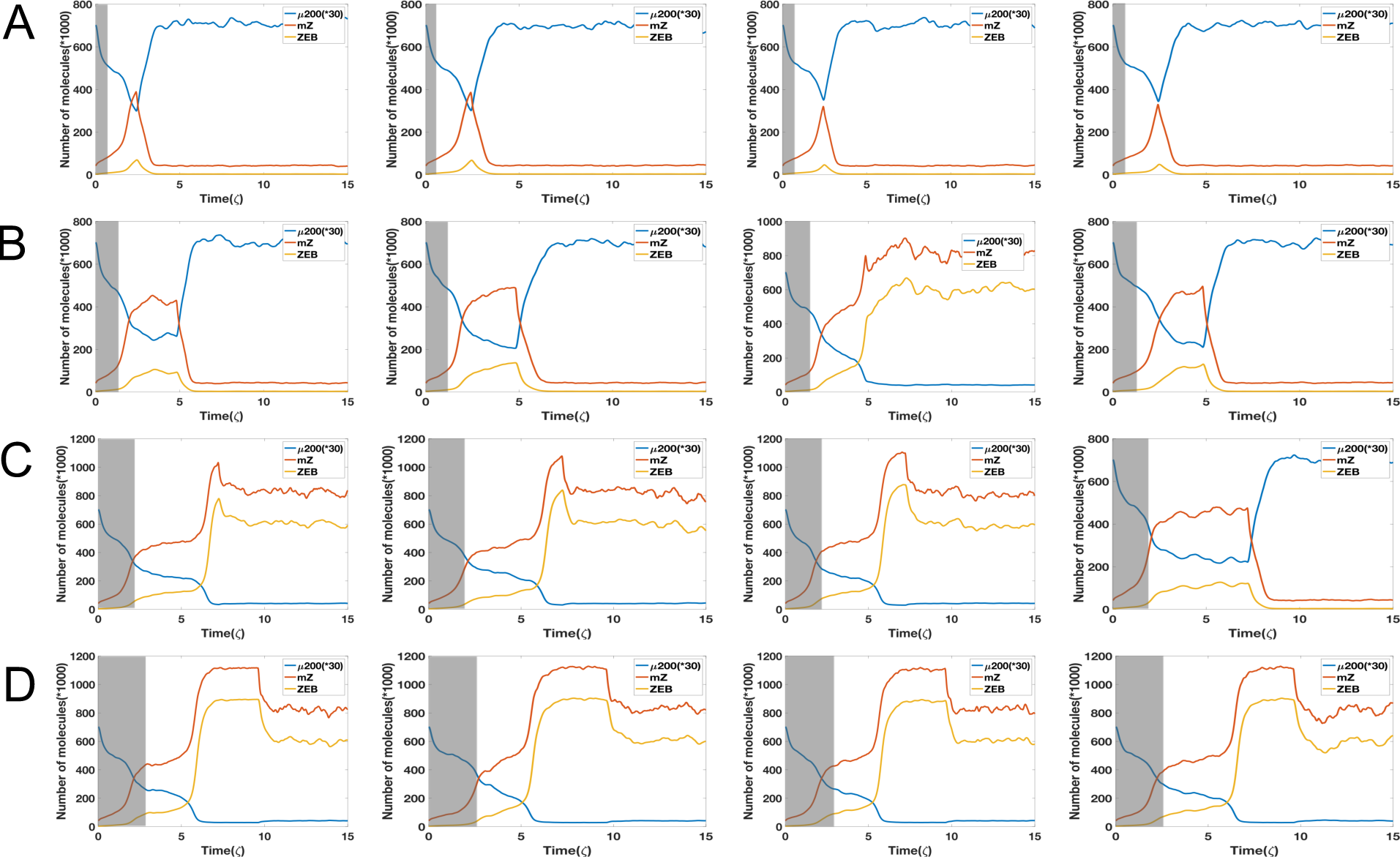
(A) Starting from I=60 K molecules (grey shaded area, bistable phase (E/M,M)), four independent dynamical examples by reducing signal I to 38 K molecules (bistable phase (E,M)) starting from time 2.4 *ζ*. (B) Same as Fig 4A but reducing signal I to 38 K molecules at time 4.8 *ζ*. (C) Same as Fig 4A but reducing signal I to 38 K molecules at time 7.2 *ζ*. (D) Same as Fig 4A but reducing signal I to 38 K molecules at time 9.6 *ζ*. System has a strong epigenetic feedback on ZEB’s inhibition on miR-200 (*α* = 0.2). The initial state is fully relaxed epithelial state with I=38. Here the variance of noise is 25 (K molecules)^2^.

**Figure 5.**
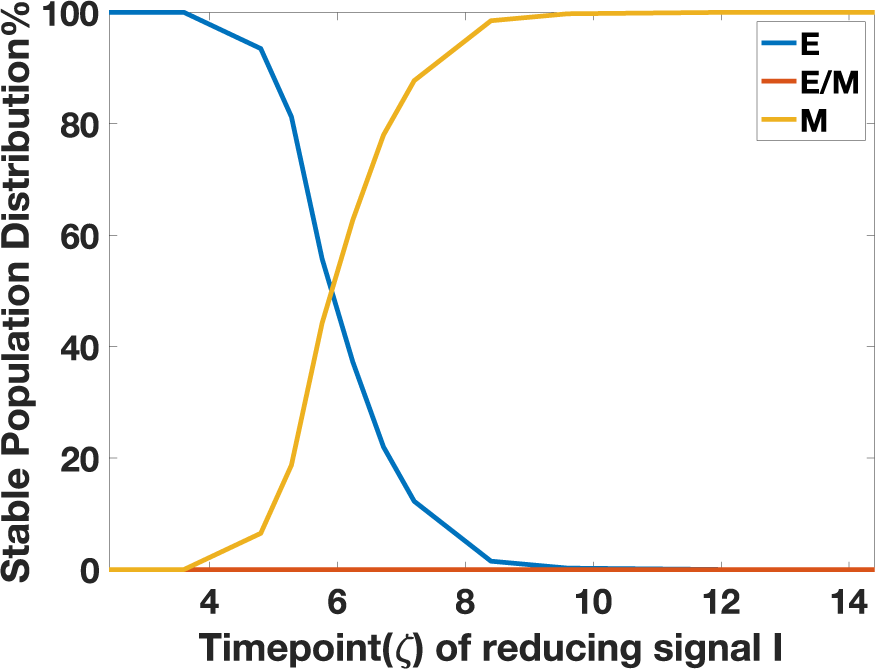
Steady state population distribution in E, E/M, M phenotypes for 1000 independent dynamical simulations of each case, as a function of the varying time durations of high I that the cells are exposed to.

### Experiments reveal the irreversibility of EMT upon prolonged EMT induction

To directly experimentally test the prediction from our model that a prolonged exposure to TGF-β allows carcinoma cells to maintain EMT characteristics even after the removal of the signal, we co-transduced a Z-CAD dual sensor system into a non-tumorigenic epithelial cell line MCF10A (Fig. 6A). This system is comprised of two individual sensors: the first component is destabilized green fluorescent protein (GFP) reporter regulated by the ZEB1 3’ untranslated regions (UTR) and the second component is red fluorescent protein (RFP) reporter driven by E-cadherin promoter. As discussed above, ZEB1 forms a double-negative feedback loop with miR-200 family, which induces MET by binding ZEB1 3’UTR, thus suppressing ZEB1 translation (35,36,37). On the other hand, the loss of E-Cadherin is hallmark of EMT; it is typically mediated via direct transcriptional suppression by ZEB1 and other master transcriptional factors that target E-Boxes presented in the E-cadherin promoter (38).

**Figure 6.**
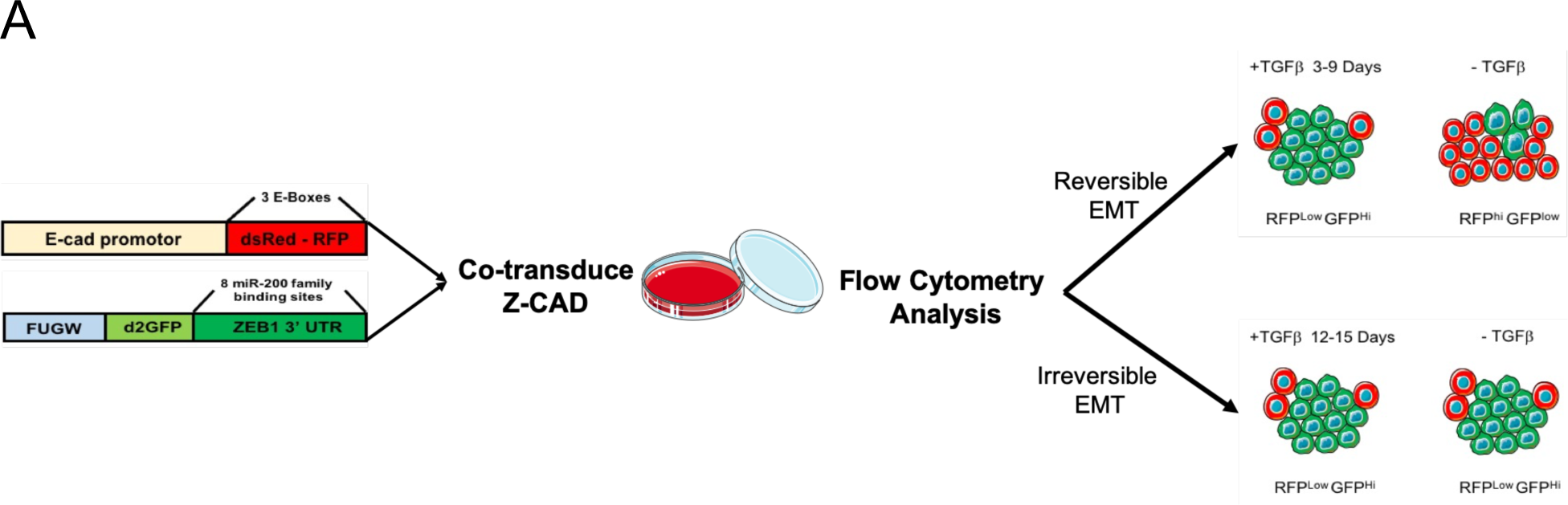
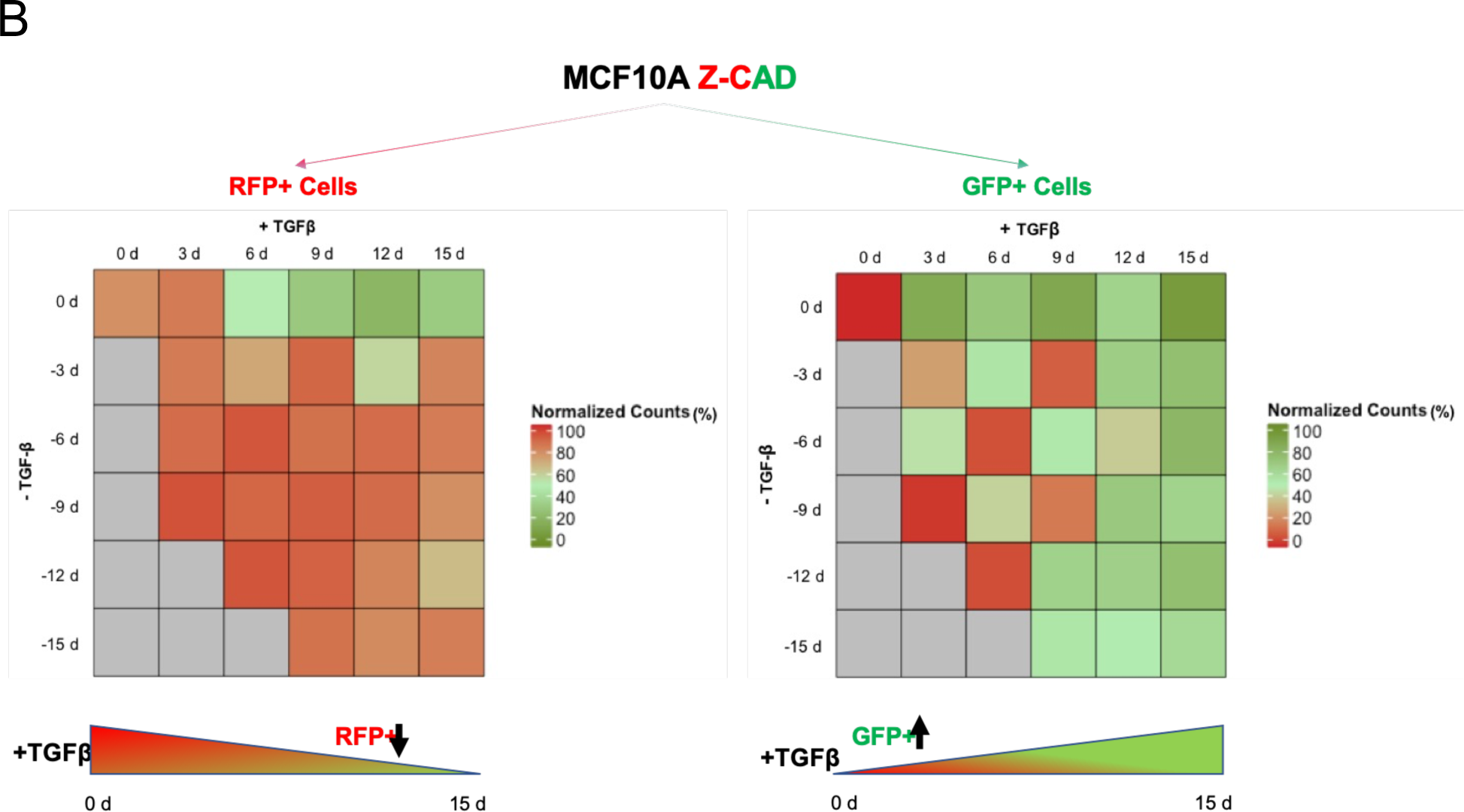
TGFβ1-treated MCF10A breast cancer cells harboring the Z-cad sensor. (A) Schematic representation of the experimental design to assess the EMT induction and reversibility. The cells were predominantly RFP ^High^ at time 0 day. A transition to GFP ^high^ (ZEB1) positivity marked the onset of EMT. (B) Dynamic transition of Z-CAD cells and their reversibility on TGFβ withdrawal. TGFβ withdrawal analysis in RFP+ (E-CAD) and GFP+ (ZEB1) population to assess the EMT reversibility (n = 3 biological replicates). % population comprises both single-color positive cells (either epithelial or mesenchymal) as well as hybrid E/M cells (harboring both epithelial as well as mesenchymal traits)

We exposed MCF10A cells to varying durations of TGF-β and counted the number of GFP+ and RFP+ cells. MCF10A cells exposed to TGF-β showed an increasingly mesenchymal morphology (SI section 11). For cells exposed to TGF-β for a shorter duration (3-6 days), the cells reverted to being epithelial within a similar timeframe (3-6 days). However, for cells exposed to TGF-β for a longer duration (12-15 days), they underwent a stronger degree of EMT, as apparent from the loss of E-cadherin and increased ZEB1 expression, but did not all revert to being epithelial even after TGF-β removal for another 15 days (Fig. 6B). They maintained their changed morphology (SI section 11), increased expression of EMT transcription factor ZEB1 and decreased expression of E-cadherin.

These data illustrate that the long-term treatment of epithelial cells with TGF-β can confer a non-reversible mesenchymal phenotype, i.e. cells can maintain a mesenchymal phenotype without the need of any externally supplied TGF-β.

## Discussion

Many studies have been devoted to understanding the multiple regulatory layers underlying EMT (3). Recent computational models have indicated that the multistability of EMT is controlled by the miR-200/ZEB and miR-34/SNAIL loops (10, 22, 39, 40). Lu *et al.* have argued specifically that the tristablility is controlled by the miR-200/ZEB unit, while the miR-34/SNAIL unit functions as a noise-buffering integrator (10). A more pronounced role of ZEB1 in initiating and sustaining EMT, as compared to other EMT inducers, has been proposed experimentally as well (34).

Here, we used a phenomenological approach to investigate how various epigenetic processes can alter the dynamics of miR-200/ZEB loop, and hence the propensity of cells to undergo EMT/MET. We found that the effects of epigenetic feedback on the self-activation of ZEB are negligible, while the epigenetic feedback on the inhibition on miR-200 by ZEB can significantly increase the stability of the mesenchymal state. It becomes therefore possible for cells to stay much longer in the mesenchymal state if this feedback is strong enough to strongly suppress transitions out of this state. This behavior greatly increases the expected degree of hysteresis in EMT as was predicted earlier through mathematical models (10), and has been recently experimentally demonstrated (18,19).

Based on these results, we designed an *in silico* experiment to demonstrate the effects of epigenetic feedback regulation and ran simulations to obtain a prediction for the experiment results. Our model predicted that if the external inducing EMT signal is applied long enough and then removed, some mesenchymal cells cannot undergo the reverse process (MET) in any reasonable timeframe, suggesting they are trapped in a mesenchymal attractor (41). Besides our preliminary experimental data, this prediction also finds support in recent studies showing that a chronic exposure to TGF-β drives a stabilized EMT (34, 42). However, it is possible for cell to return to its initial state for short time treatment with an EMT-inducing signal such as TGF-*β*. Our basic assumption is that the relevant time scale governing these processes should be of the same order as those governing EMT (perhaps 5 days or so) and dynamical changes pertaining to longer periods of time are more likely to be of epigenetic origin. Of course, one could test this hypothesis directly by investigating the effects of modifying various epigenetic mechanisms by manipulating the responsible enzymatic pathways. As we investigate our experimental system more fully, we hope to report on these tests in future.

Besides epigenetics, other possible dynamical mechanisms can give rise to major differences in MET as compared to EMT. One such mechanism is secretion of TGF-β by cells in mesenchymal state that could stabilize EMT in both autocrine and paracrine ways (34, 43), but also makes estimating the population dynamics more complex as EMT is no longer a completely cell-autonomous process. This type of stabilization could also occur via other cell-signaling processes such as those mediated by the Notch-Jagged pathway (44). Another hypothesis claims that EMT and MET trajectories are asymmetric without necessarily invoking epigenetic differences, i.e. when a cell undergoes MET, it reaches a new epithelial state that is different from the initial epithelial state (19). Again, the expected time scale for these events is shorter and hence changes that require long-term exposure are likely to have epigenetic components.

There are increasing indications that cells that have undergone some form of EMT can be crucial for enabling tumors to resist treatment, disperse to distant locations, and eventually start growing secondary lesions. Our results indicate that epigenetic processes may play an important role in stabilizing cells that have been induced to undergo EMT by some seminal event but may no longer be subjected to the same micro-environmental factors.

