## Supplementary Information for "A possible role for epigenetic feedback regulation in the dynamics of the Epithelial-Mesenchymal Transition (EMT)"

#### 1.Theoretical model for EMT

In this EMT network, microRNA, mRNA and protein affect each other via different mechanisms. According to the framework built by Lu. et al (10), the deterministic equations for miR-200/ZEB circuit with the external signal as SNAIL are:

$$\dot{\mu}_{200} = g_{\mu_{200}} H^s(Z, \lambda_{z,\mu_{200}}) H^s(S, \lambda_{S,\mu_{200}}) - m_z Y_\mu(\mu_{200}) - k_\mu \mu_{200}$$

$$\dot{m}_z = g_{m_z} H^s(Z, \lambda_{z,m_z}) H^s(S, \lambda_{s,m_z}) - m_z Y_m(\mu_{200}) - k_{m_z} m_z$$

$$\dot{Z} = g_z m_Z L(\mu_{200}) - k_z Z$$

and those for miR-34/SNAIL circuit with I as an external signal are:

$$\dot{\mu}_{34} = g_{\mu_{34}} H^s(S, \lambda_{s,\mu_{34}}) - m_s Y_\mu(\mu_{34}) - k_{\mu_{34}} \mu_{34}$$

$$\dot{m}_s = g_{m_s} H^s(S, \lambda_{s,m_s}) H^s(I, \lambda_{I,m_s}) - m_s Y_m(\mu_{34}) - k_{m_s} m_s$$

$\dot{S} = g_s m_s L(\mu_{34}) - k_s S$

So, the combined circuit is driven by I is given by:

$\dot{\mu}_{200} = g_{\mu_{200}} H^s(Z, \lambda_{z,\mu_{200}}) H^s(S, \lambda_{S,\mu_{200}}) - m_z Y_\mu(\mu_{200}) - k_\mu \mu_{200}$

$\dot{m}_z = g_{m_z} H^s(Z, \lambda_{z,m_z}) H^s(S, \lambda_{s,m_z}) - m_z Y_m(\mu_{200}) - k_{m_z} m_z$

$\dot{Z} = g_z m_Z L(\mu_{200}) - k_z Z$

$\dot{\mu}_{34} = g_{\mu_{34}} H^s(S, \lambda_{s,\mu_{34}}) H^s(Z, \lambda_{z,\mu_{34}}) - m_s Y_\mu(\mu_{34}) - k_{\mu_{34}} \mu_{34}$

$\dot{m}_s = g_{m_s} H^s(S, \lambda_{s,m_s}) H^s(I, \lambda_{I,m_s}) - m_s Y_m(\mu_{34}) - k_{m_s} m_s$

$\dot{S} = g_s m_s L(\mu_{34}) - k_s S$

where $g$ is the innate synthesis rate for corresponding microRNA/mRNA/protein, $k$ is the corresponding innate degradation rate. Here $H^S$ represents the shifted Hill function which is defined as:

$$H^s(B) = \frac{1 + \lambda \left(\frac{B}{B_0}\right)^{n_B}}{1 + \left(\frac{B}{B_0}\right)^{n_B}}$$

where $\lambda$ is the fold change regulated by protein B. $\lambda$>1 for activation and $\lambda$<1 for inhibition. The function $Y$ represents degradation of microRNA or mRNA due to microRNA-mRNA binding ($n$ is the number of binding sites). The function $L$ represents translational inhibition. They can be written as:

$$L(\mu) = \sum_{i=0}^{n} l_i \, C_n^i M_n^i(\mu)$$

$$Y_m(\mu) = \sum_{i=0}^{n} \gamma_{mi}\, C_n^i M_n^i(\mu)$$

$$Y_\mu(\mu) = \sum_{i=0}^{n} i\gamma_{\mu i}\, C_n^i M_n^i(\mu)$$

where

$$M_n^i(\mu) = \frac{\left(\frac{\mu}{\mu_0}\right)^i}{\left(1 + \frac{\mu}{\mu_0}\right)^n}$$

$$\sum_{i=0}^{n} C_n^i M_n^i(\mu) = 1$$

Here $l_i, \gamma_{mi}, \gamma_{\mu i}$ correspond to the individual translation rate of mRNA, individual degradation rate for mRNA and microRNA respectively. All details of microRNA-mediated regulation can be found in Lu et al. (1).

### 2.Parameters for the EMT model

**Table SI 1. List of parameters used in shifted Hill functions**

| Description | Fold change | Value | # of binding sites | Value | Threshold | Value (K molecules) |
|---|---|---|---|---|---|---|
| Inhibition on miR-200 by ZEB | $\lambda_{Z,\mu_{200}}$ | 0.1 | $n_{Z,\mu_{200}}$ | 3 | $Z^0_{\mu_{200}}$ | 220 |
| Inhibition on miR-200 by SNAIL | $\lambda_{S,\mu_{200}}$ | 0.1 | $n_{S,\mu_{200}}$ | 2 | $S^0_{\mu_{200}}$ | 180 |

| | | | | | | |
|---|---|---|---|---|---|---|
| Self-activation of ZEB | $\lambda_{Z,m_z}$ | 7.5 | $n_{Z,m_z}$ | 2 | $Z^0_{m_z}$ | 25 |
| Activation on ZEB by SNAIL | $\lambda_{S,m_z}$ | 10.0 | $n_{S,m_z}$ | 2 | $S^0_{m_z}$ | 180 |
| Inhibition on miR-34 by SNAIL | $\lambda_{S,\mu_{34}}$ | 0.1 | $n_{S,\mu_{34}}$ | 1 | $S^0_{\mu_{34}}$ | 300 |
| Inhibition on miR-34 by ZEB | $\lambda_{Z,\mu_{34}}$ | 0.2 | $n_{Z,\mu_{34}}$ | 2 | $Z^0_{\mu_{34}}$ | 600 |
| Self-inhibition of SNAIL | $\lambda_{S,m_s}$ | 0.1 | $n_{S,m_s}$ | 1 | $S^0_{m_s}$ | 200 |
| Activation on SNAIL by external signal I | $\lambda_{I,m_s}$ | 10 | $n_{I,m_s}$ | 2 | $I^0_{m_s}$ | 50 |

**Table SI 2. List of parameters for function *Y* and *L*.**

| n (# of miRNA binding sites) | 0 | 1 | 2 | 3 | 4 | 5 | 6 |
|---|---|---|---|---|---|---|---|
| $l_i$(hour$^{-1}$) | 1 | 0.6 | 0.3 | 0.1 | 0.05 | 0.05 | 0.05 |
| $\gamma_{mi}$(hour$^{-1}$) | 0 | 0.04 | 0.2 | 1 | 1 | 1 | 1 |
| $\gamma_{\mu i}$(hour$^{-1}$) | 0 | 0.005 | 0.05 | 0.5 | 0.5 | 0.5 | 0.5 |
| $n_{\mu_{200}}$ | 6 | $n_{\mu_{34}}$ | | | | | 2 |
| $\mu^0_{200}$ | 10K | $\mu^0_{34}$ | | | | | 10K |

**Table SI 3. List of other parameters used in EMT model.**

| Synthesis rate | Value (molecules/hour) | Degradation rate | Value ($hour^{-1}$) | Translation rate | Value ($hour^{-1}$) |
|---|---|---|---|---|---|
| $g_{\mu_{200}}$ | 2.1K | $k_{\mu_{200}}$ | 0.05 | $g_z$ | 0.1K |
| $g_{m_z}$ | 11 | $k_{m_z}$ | 0.5 | $g_s$ | 0.1K |
| $g_{\mu_{34}}$ | 1.35K | $k_z$ | 0.1 | | |
| $g_{m_s}$ | 90 | $k_{\mu_{34}}$ | 0.05 | | |
| | | $k_{m_s}$ | 0.5 | | |
| | | $k_s$ | 0.125 | | |

**3.External signal noise on SNAIL**

The external signal I that we use here can be written as the stochastic differential equation:

$$\dot{I} = \beta(I_0 - I) + \eta(t)$$

where $\eta(t)$ satisfies the condition that $< \eta(t), n(t') \geq \Gamma\delta(t - t')$. Here $I_0$ is set at 50 K molecules, $\beta$ as 0.04 $hour^{-1}$, and Γ as 50 $(K\ molecules/hour)^2$.

The initial value of I is fixed to lie at the middle of the tristable region (E, E/M, M).

**4. Epigenetic feedback regulation term**

In the EMT model, we tested epigenetic feedback through two different pathways. The dynamic equation of epigenetic feedback on ZEB's self-activation is:

$$\dot{Z}^0_{m_z} = \frac{Z^0_{m_z}(0) - Z^0_{m_z} - \alpha Z}{\zeta}$$

Simialry, epigenetic feedback on ZEB's inhibition on miR-200 is modeled via:

$$\dot{Z}^0_{\mu_{200}} = \frac{Z^0_{\mu_{200}}(0) - Z^0_{\mu_{200}} - \alpha Z}{\zeta}$$

where $\zeta$ is a timescale factor and chosen to be 100 (hours). $\alpha$ represents the strength of epigenetic feedback. Larger $\alpha$ corresponds to stronger epigenetic feedback. $\alpha$ has an upper bound because of the restriction that the numbers of all molecules must be positive. For ZEB's self-activation, high level of ZEB can activate the expression of ZEB itself due to this epigenetic regulation. Meanwhile, for ZEB's inhibition on miR-200, high levels of ZEB can suppress the synthesis of miR-200.

In our EMT model, we used $\zeta$ = 100 hours as the unit of time, because the timescale in our feedback-dependent simulations depends on not only the noise, but also the value of $\zeta$.

**5. Simple model for understanding EMT: The SATS model**

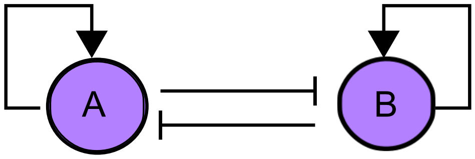


**Figure S1.** The regulatory network of self-activating toggle switch----SATS (33).

To gain more confidence in our results regarding the EMT circuit, we begin with a simpler case – the self-activating toggle switch (SATS). A SATS consists of two mutually inhibiting transcription factors (TFs) and has two states – 'A' state (A high, B low), and 'B' state (A low, B high) (Fig S1). The dynamics of a SATS is given by:

$$\frac{dA}{dt} = g_A H^{S_{AA}}(A, \lambda_{AA}, n_{AA}, A_A^0) H^{BA}(B, \lambda_{BA}, n_{BA}, B_A^0) - k_A A$$

$$\frac{dB}{dt} = g_B H^{S_{BB}}(B, \lambda_{BB}, B_B^0) H^{AB}(A, \lambda_{AB}, n_{AB}, A_B^0) - k_B B$$

The epigenetic feedback in SATS can be represented by:

$$\dot{A_A^0} = \frac{(A_A^0(0) - A_A^0 - \alpha A)}{\zeta} \qquad (*)$$

$$\dot{B_A^0} = \frac{(B_A^0(0) - B_A^0 - \alpha B)}{\zeta}$$

We studied two cases:1. feedback on the self-activation of A; 2. feedback on the inhibition on A by B. The term $\alpha \cdot A$ or $\alpha \cdot B$ represents the epigenetic feedback. Because of this epigenetic feedback, for example in equation (*), if A is expressed, the threshold decreases, which finally causes A to be expressed at a higher level. Here, $\alpha$ has maximum values due to the minus sign (i.e. the threshold can not be negative), and for each case, the maximum value of $\alpha$ is different.

**6.Methods used in SATS model study**

**ODE simulations**

Here we added a Gaussian white noise term to the dynamic equations to trigger transitions between the two states. When we started from all cells in A state, we can observe the transitions by simply using the Euler method. We simulated this for 1000 times, counted the number of trajectories leading to state A and state B for each time point, and calculated the percentage of these two states. By varying $\alpha$, we can see how the population distribution changes. Increasing $\alpha$ means a stronger epigenetic feedback.

**Stochastic method**

The chemical rate equations for a SATS model:

$$\frac{dA}{dt} = -r_{AB}A + r_{BA}B$$

$$\frac{dB}{dt} = -r_{BA}B + r_{AB}A$$

And the corresponding solution is:

$$A(t) = \frac{r_{BA} + r_{AB}e^{-(r_{AB}+r_{BA})t}}{r_{AB} + r_{BA}}$$

By fitting the population distribution curve, we can get the fitting values of $r_{AB}$ and $r_{BA}$. According to the Gillespie method, here we can generate two random numbers to determine when the next transition would happen and which one (A to B or B to A). Given a constant time, we can count the number of transitions, and plot it as a function of $\alpha$.

### 7.Results of SATS model

#### Epigenetic feedback on A's self-activation

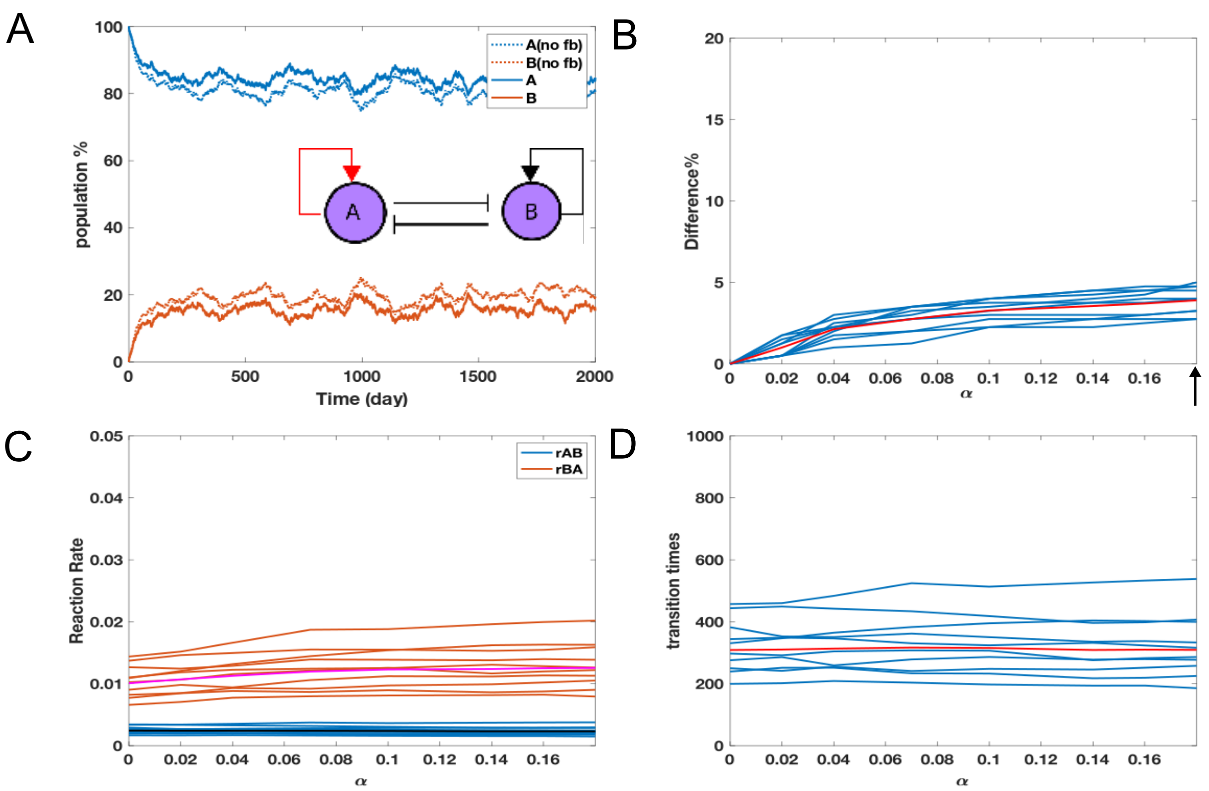


**Figure S2.** (A) A sample showing the population change as a function of time, for epigenetic feedback added to self-activation of transcription factor A. The percentage is calculated based on 1000 independent simulations. Dashed lines represent no epigenetic feedback case ($\alpha = 0$), and solid lines are with feedback ($\alpha$ value is marked by arrow in Fig. S2(B), $\alpha = 0.18$). Here, the timescale is dynamic timescale. (B) The difference between the frequency of solutions converging to the (A high, B low) state, as a function of $\alpha$. (C) Chemical reaction rates as a function of $\alpha$. (D) Transition times as a function of $\alpha$ (from Gillespie method). In all the three figures here, same simulation was repeated 10 times (trajectories plotted here to quantify the error, and the different color in each plot represents the average result.

When the epigenetic feedback is on the self-activation of A, it does not largely change the steady state distribution of the system. The reaction rates as well as the transition times during same time period remain almost constant, even if the feedback is very strong.

### Epigenetic feedback on B's inhibition on A

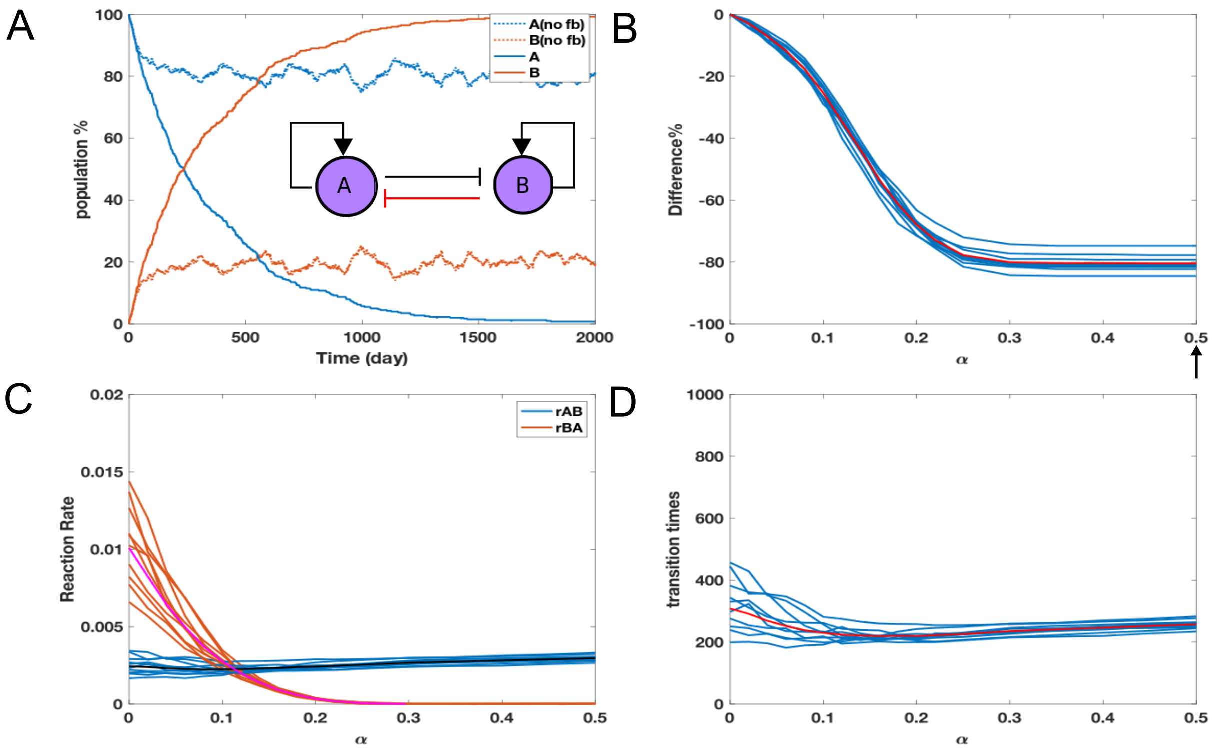


**Figure S3.** Similar data analysis method as shown in Fig. S2, but for epigenetic feedback added to the inhibition of . (A) A sample showing the population change as a function of time. The percentage is calculated based on 1000 times independent simulations. Dashed lines represent no epigenetic feedback case ($\alpha = 0$), and solid lines are with feedback ($\alpha$ value is marked by arrow in SI 3(B), $\alpha = 0.5$). (B) The difference between A's distribution population as a function of $\alpha$. (C) Chemical reaction rates as a function of $\alpha$. After reaching certain point, $r_{AB}$ (rate from state A to B) <$r_{BA}$ (rate from state B to A). (D)Transition times as a function of $\alpha$ (from Gillespie method).

When the epigenetic feedback in incorporated in the inhibtion of A by B, the equilibrium population distribution tends to move towards a higher percentage of cells in (B high, A low) state as compared to that in (A high, B low) state. This shift can be understood as following: a stronger inhibition of B on A would prevent the cells which are already in (B high, A low) state from transitioning to (A high, B low) state. Asymptotically, when the feedback is strong enough, all cells will be in the B state. From the perspective of reaction rates, this epigenetic feedback would significantly reduce the transition rate from (B high, A low) to (A high, B low).

The results of SATS model indicate that when the epigenetic feedback is on the self-activation of the TF, the system remains almost unchanged. But when this feedback is on the mutual inhibition between two TFs or innate transcription rate, the system would change with the strength of the feedback and show some “stabilized” states.

**8.Parameters used for SATS model**

**Table SI 4. List of parameters in SATS model**

| | Production rate | | Value | Degradation rate | | Value |
|---|---|---|---|---|---|---|
| TF A | $g_A$ | | 5 | $k_A$ | | 0.1 |
| TF B | $g_B$ | | 5 | $k_B$ | | 0.1 |
| | Fold change | Value | # of binding sites | Value | Threshold | Molecules |
| Inhibition of B by A | $\lambda_{A,B}$ | 0.1 | $n_{A,B}$ | 1 | $A_B^0$ | 120 |
| Inhibition of A by B | $\lambda_{B,A}$ | 0.1 | $n_{B,A}$ | 1 | $B_A^0$ | 120 |
| Self-activation of A | $\lambda_{A,A}$ | 10 | $n_{A,A}$ | 4 | $A_A^0$ | 80 |
| Self-activation of B | $\lambda_{B,B}$ | 10 | $n_{B,B}$ | 4 | $B_B^0$ | 80 |
| External signal S=600 | | | | | | |
| $n_{S,A} = n_{S,B} = 1, S_A^0 = S_B^0 = 500, \lambda_{S,A} = \lambda_{S,B} = 1$ | | | | | | |

**9.The effects of noise**

In order to test the effects of external signal noise in EMT model, we tried 10 different values of standard deviation for a given initial condition and analyzed the results. The initial condition is that 100 % cells are in a M state and there is strong epigenetic feedback on ZEB's inhibition on miR-200 ($\alpha = 0.2$). The mean value of I used here is 51.3 K molecules, which corresponds to tristable phase {E, E/M, M} (Fig S4).

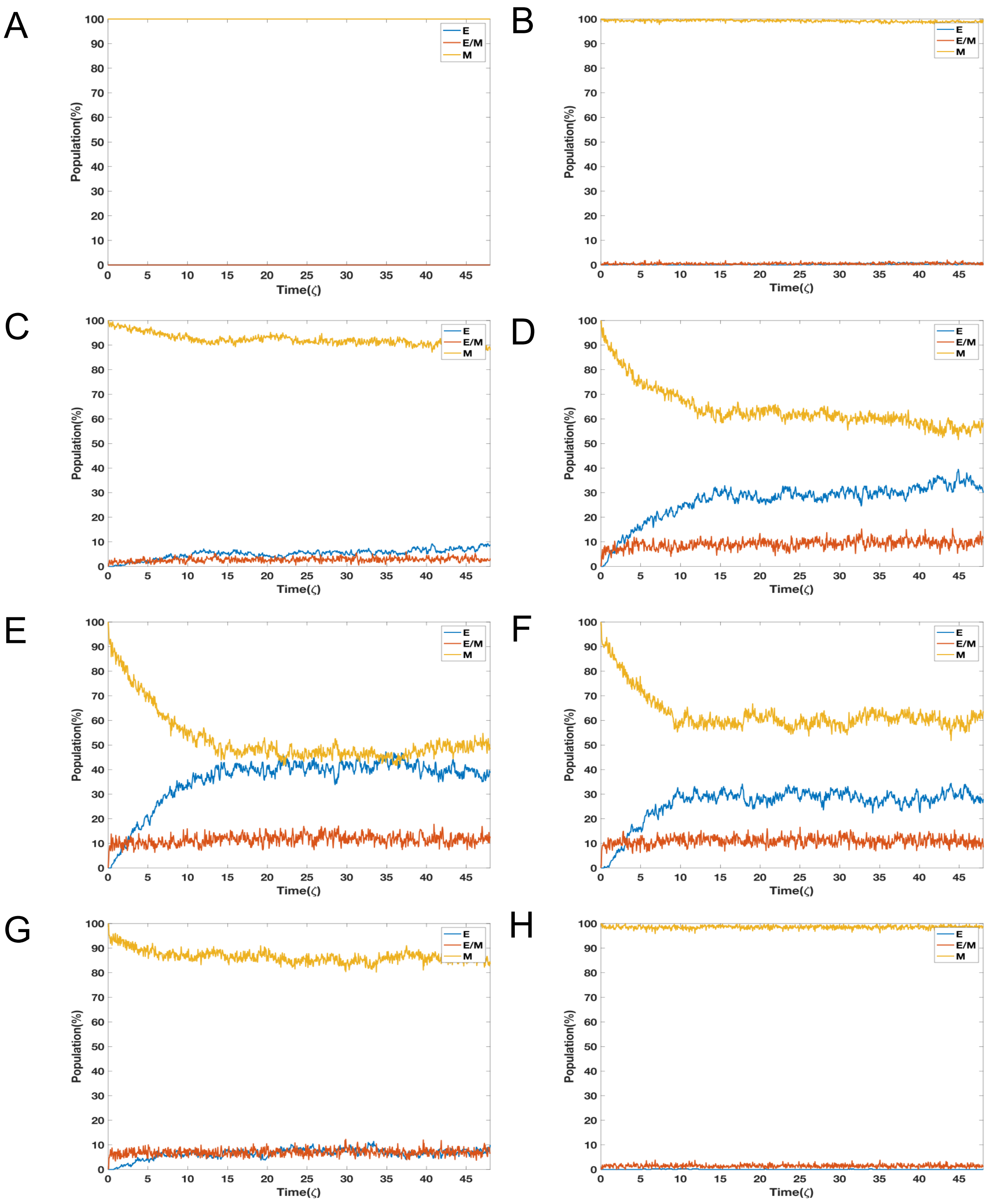


**Figure S4.** Population distribution results. (A) The standard deviation $\sigma < 30$. (B) $\sigma = 40$. (C) $\sigma = 50$. (D) $\sigma = 70$. (E) $\sigma = 100$. (F) $\sigma = 120$. (G) $\sigma = 150$. (H) $\sigma = 200$.

Starting from the M state for low standard variation ($\sigma < 50$) case, no transition is observed (Fig. S4A, B). When standard deviation is too large, a large percentage of cells maintain themselves in an M state (Fig. S4G,H) because….. <Also comment on Fig S4C-F>. From these results, if the noise is mostly in tristable region, it's not enough to trigger the transition. Meanwhile, the magnitude of noise would affect the population distribution and time needed to reach it. It's kind of trade-off between these factors.

In our simulation, we chose $\gamma = 50, \tau = 0.01, \sigma = \sqrt{\frac{\gamma}{\tau}} \approx 70$, so we can observe the reasonable timescale as well as stable distribution compared with our preliminary experimental results.

**10.Experiment methods**

MCF10A cells were maintained in DMEM/F12 (Gibco) supplemented with 5 % horse serum, 20 ng/mL epidermal growth factor (EGF), 0.5 μ g/mL hydrocortisone, 5 μ g/mL insulin, 100 ng/mL cholera toxin, and antibiotic. The MCF10A cells containing the Z-CAD sensor were obtained from Dr. Jefferey Rosen (Baylor College of Medicine, Houston, TX) (45). The Z-CAD cells were treated with TGF-β (5 ng/mL) to induce EMT over the course of several days. Flow cytometry analysis were performed every 3rd Day to demonstrate the E-M transition. Importantly, we were able to identify changes over time in a transitioning population, demonstrating the ability to observe dynamic changes displaying reversible EMT characteristics. Finally, we also showed that a with prolonged TGF-β treatment, Z-CAD cells have permanently undergone EMT and are irreversible, as identified by their Z-cad sensor fluorescence pattern.

### 11.Experimental morphology results

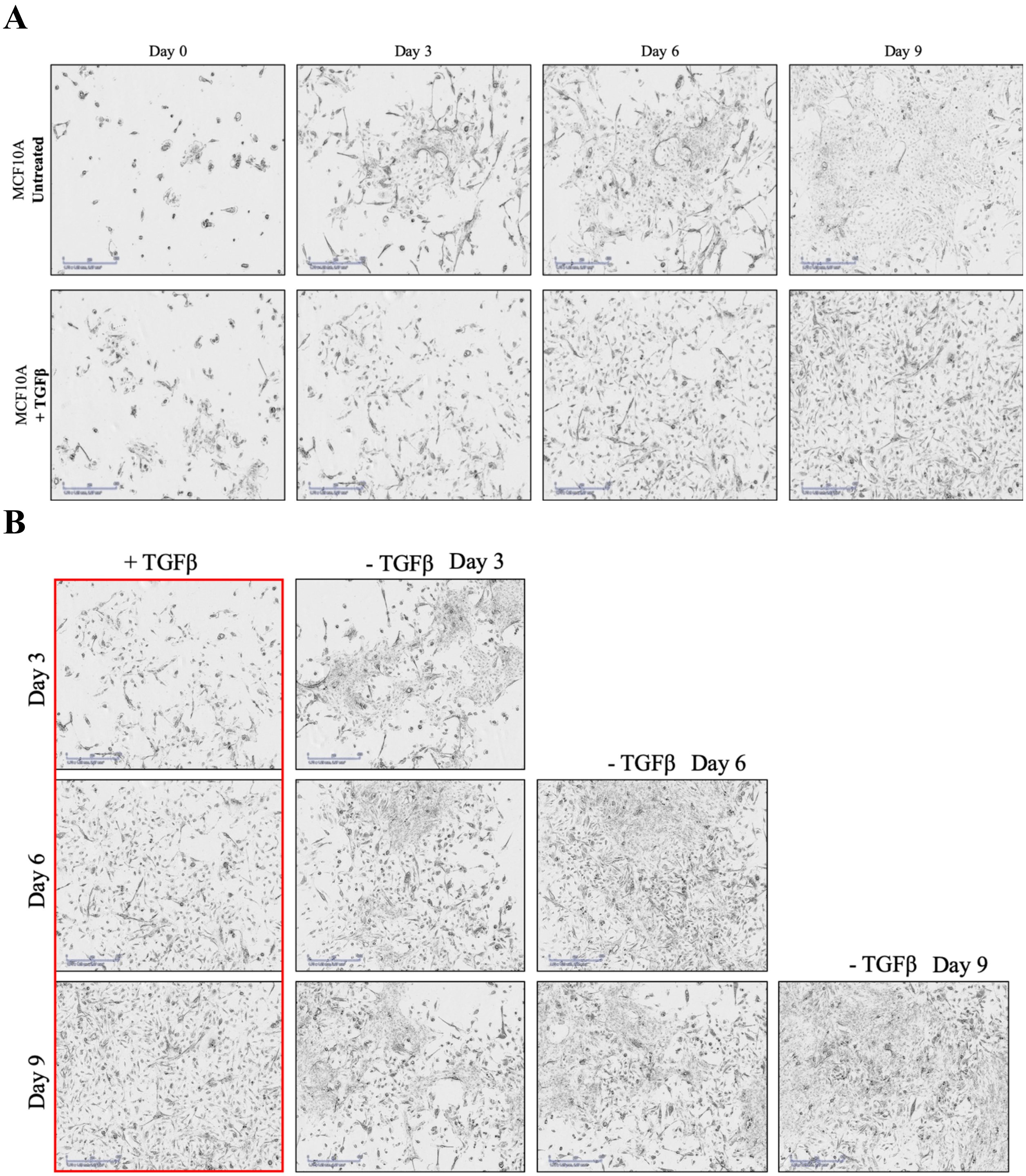


**Figure S5.** (A)Morphology pictures of TGFβ1-treated MCF10A breast cancer cells vs untreated cells. (B)Morphology pictures of TGFβ1-treated MCF10A breast cancer cells vs results after withdrawing TGFβ1 for 3-9 days.
